# Variation in the microbiome of the urogenital tract of female koalas (*Phascolarctos cinereus*) with and without ‘wet bottom’

**DOI:** 10.1101/099945

**Authors:** Alistair R. Legione, Jemima Amery-Gale, Michael Lynch, Leesa Haynes, James R. Gilkerson, Fiona M. Sansom, Joanne M. Devlin

## Abstract

Koalas (*Phascolarctos cinereus*) are iconic Australian marsupials currently threatened by several processes. Infectious reproductive tract disease, caused by *Chlamydia pecorum*, and koala retrovirus infection are considered key drivers of population decline. The clinical sign of ‘wet bottom’, a staining of the rump associated with urinary incontinence, is often caused by chlamydial urogenital tract infections. However, wet bottom has been recorded in koalas free of *C. pecorum*, suggesting other causative agents in those individuals. Current understanding of the bacterial community of the koala urogenital tract is limited. We used 16S rRNA diversity profiling to investigate the microbiome of the urogenital tract of ten female koalas. This was to produce baseline data on the female koala urogenital tract microbiome, and to undertake preliminary investigations of potential causative agents of wet bottom, other than *C. pecorum*. Five urogenital samples were processed from koalas presenting with wet bottom and five were clinically normal. We detected thirteen phyla across the ten samples, with *Firmicutes* occurring at the highest relative abundance (77.6%). The order *Lactobacillales*, within the *Firmicutes*, comprised 70.3% of the reads from all samples. After normalising reads using DESeq2 and testing for significant differences (*P* < 0.05), there were 25 operational taxonomic units (OTUs) more commonly found in one group over the other. The families *Aerococcaceae* and *Tissierellaceae* both had four significantly differentially abundant OTUs. These four *Tissierellaceae* OTUs were all significantly more abundant in koalas with wet bottom.

**Importance:** This study provides an essential foundation for future investigations of both the normal microflora of the koala urogenital tract, and better understanding of the causes of koala urogenital tract disease. Koalas in the states of Queensland and New South Wales are currently undergoing decline, and have been classified as vulnerable populations. Urogenital tract disease is a leading cause of hospital admissions in these states, yet previously little was known of the normal flora of this site. Wet bottom is a clinical sign of urogenital tract disease, which is often assumed to be caused by *C. pecorum* and treated accordingly. Our research highlights that other organisms may be causing wet bottom, and these potential aetiological agents need to be further investigated to fully address the problems this species faces.

## Introduction

The koala (*Phascolarctos cinereus*) is an iconic marsupial species endemic to Australia. Northern koala populations, in the states of Queensland and New South Wales, are currently declining due to impacts from disease and increased urbanisation. A significant pathogen of koalas, *Chlamydia pecorum*, has been a main focus of koala infectious disease investigations since its discovery. *C. pecorum* infection of the conjunctiva or urogenital tract can lead to blindness and infertility in koalas, respectively, greatly impacting population fecundity and survivability (1, 2). *C. pecorum* is commonly associated with the clinical sign known as ‘wet bottom’ or ‘dirty tail’ (3). This staining or scalding of the rump is associated with cystitis due to *C. pecorum* infection in some populations (4), but recently samples from a large number of koalas from Victorian populations with mild wet bottom were negative via qPCR for *C. pecorum* (5). In particular, koalas in a population considered at the time to be free of *C. pecorum* (6) had a similar prevalence and severity of wet bottom to populations where *C. pecorum* occurred in more than 35% of koalas tested. Further analysis demonstrated that whilst wet bottom could be significantly linked to the detection of *C. pecorum* infection in male Victorian koalas, this relationship did not exist in females (7). It may be that an unidentified organism is causing these mild clinical signs of disease in koalas. To date there has not been extensive research to determine the normal flora of the koala urogenital tract, making it difficult to use traditional microbiological techniques to determine species of interest. Modern sequencing technology, specifically 16S rRNA biodiversity profiling, was used to improve our understanding of the microbiome of the urogenital tract of koalas, and to undertake preliminary comparisons of the microbiome of female koalas with and without mild wet bottom.

## Results

### Clinical status of koalas

Urogenital samples previously collected from ten koalas as a component of population health monitoring were selected from an archive of samples available at our institute (7, 8). The criteria for selection was based on adequate cold storage of samples in an appropriate buffer. Five samples that met our criteria, taken from koalas with wet bottom, were available. An additional five samples, taken from koalas with no clinical signs of disease, were selected from the same population. Of the five koalas with wet bottom, the median wet bottom clinical score was 3 (ranging from 2 – 4). The five clinically healthy animals all had wet bottom clinical scores of 0. All koalas were negative for *Chlamydiaceae* using a pan-*Chlamydiaceae* qPCR.

### Analysis and processing of sequencing data

A total of 2,295,607 paired reads were obtained across the ten samples, ranging between 189,315 to 312,131 reads per sample. The GC content of the reads was 51.8%. Merging paired reads, trimming 5’ and 3’ ends, quality filtering to remove errors and discarding merged sequences shorter than 400 bp resulted in a total of 1,347,512 reads. Dereplication resulted in 275,642 unique reads for clustering into operational taxonomic units (OTUs). Through the clustering process, it was determined that 3953 unique reads were chimeric,representing 24,376 filtered reads. The non-chimeric unique reads were clustered into 261 OTUs, 7 of which were either chloroplasts or mitochondria and were subsequently removed from the analysis. In total 1,946,587 reads, from 2,221,529 merged reads (87.6%) were matched to the clustered OTUs. Within samples, this ranged from 162,343 (82% of available reads) to 254,327 (92.1% of available reads) (Table 1). For comparison, the same filtering and clustering methodology was run without the removal of singletons, which resulted in the clustering of reads into 592 OTUs.

**Table 1.** Koala wet bottom score, read metrics and relative abundance data from ten samples submitted for 16S rRNA amplicon sequencing. All koalas were female and sampled from French Island, Victoria, Australia in 2011.

| Koala/Sample name | K1 | K2 | K3 | K4 | K5 | K31 | K49 | K55 | K59 | K70 |
| --- | --- | --- | --- | --- | --- | --- | --- | --- | --- | --- |
| **Wet bottom score\*** | 0 | 0 | 0 | 0 | 0 | 2 | 3 | 3 | 4 | 3 |
| **Merged reads** | 253256 | 211620 | 186912 | 220410 | 185592 | 183126 | 199985 | 263685 | 216495 | 300448 |
| **Reads after filtering** | 156100 | 134940 | 118418 | 132125 | 112823 | 110292 | 116321 | 160328 | 136996 | 169169 |
| **Reads clustered to OTUs** | 225868 | 178678 | 169576 | 203062 | 166906 | 162343 | 177452 | 216270 | 192105 | 254327 |
| **Absolute OTUs** | 93 | 66 | 86 | 89 | 74 | 55 | 61 | 74 | 76 | 126 |
| **Standardised OTUs<sup>^</sup> ± SD<sup>+</sup>** | 88.8 ± 1.7 | 64.1 ± 1.2 | 85.4 ± 0.7 | 88 ± 0.9 | 73.7 ± 0.6 | 54.9 ± 0.3 | 59.2 ± 1.4 | 69.2 ± 1.9 | 72.9 ± 1.5 | 123.4 ± 1.3 |
| **Phyla<sup>#</sup>** |  |  |  |  |  |  |  |  |  |  |
| *Acidobacteria* | - | - | - | - | < 0.01% | - | - | - | - | 0.01% |
| *Actinobacteria* | 5.47% | 9.06% | 2.92% | 0.17% | 0.03% | 3.27% | 0.66% | 1.50% | 0.30% | 0.19% |
| *Armatimonadetes* | < 0.01% | < 0.01% | - | - | < 0.01% | - | - | - | - | - |
| *Bacteroidetes* | 0.57% | 0.05% | 2.14% | 1.72% | 0.21% | 0.33% | 0.05% | 9.05% | 1.00% | 50.53% |
| *Cyanobacteria* | < 0.01% | - | < 0.01% | - | - | - | - | - | - | 0.02% |
| *Deferribacteres* | - | - | - | - | - | - | - | - | - | < 0.01% |
| *Firmicutes* | 92.92% | 89.57% | 85.67% | 79.17% | 98.92% | 80.35% | 40.92% | 84.88% | 95.65% | 39.09% |
| *Fusobacteria* | 0.02% | < 0.01% | < 0.01% | 0.07% | < 0.01% | < 0.01% | - | < 0.01% | 0.02% | 1.09% |
| *Planctomycetes* | - | - | < 0.01% | - | 0.01% | - | - | - | < 0.01% | 0.80% |
| *Proteobacteria* | 0.24% | 0.15% | 1.66% | 1.51% | 0.45% | 0.23% | 56.90% | 0.19% | 2.37% | 2.70% |
| *Synergistetes* | 0.08% | 0.02% | 0.30% | 0.31% | 0.01% | - | - | < 0.01% | 0.02% | 4.35% |
| *TM7* | 0.02% | 0.50% | 0.21% | - | < 0.01% | 1.38% | 0.05% | 2.86% | < 0.01% | 0.02% |
| *Verrucomicrobia* | < 0.01% | < 0.01% | < 0.01% | - | 0.02% | < 0.01% | - | - | 0.01% | 0.69% |
| **Unassigned** | 0.69% | 0.65% | 7.07% | 17.04% | 0.34% | 14.44% | 1.42% | 1.52% | 0.61% | 0.52% |

\* Wet bottom score ranges from 0 (absent) to 10 (most severe) (37)
<sup>^</sup> The average number of OTUs detected in 100 iterations of subsampling to a depth of 160,000 reads
<sup>+</sup> Standard deviation
<sup>#</sup> Phyla assigned using QIIME (40) script **assign\_taxonomy.py** utilising Greengenes (42) curated 16S rRNA library

### Phylum presence and relative abundance

In total, 13 phyla were detected in the ten samples (Table 1), with Firmicutes occurring at the highest relative abundance (77.61%). Just over a third of the OTUs were classified as Firmicutes (95/254), followed by Proteobacteria (59/254) and the Bacteroidetes (35/254). When samples were split into the two groups, koalas without wet bottom had 89.3% of reads classified as Firmicutes, followed by OTUs which could not be assigned using the 90% similarity threshold (5.2%) and Actinobacteria (3.5%). Koalas with wet bottom had 68.2% reads assigned to OTUs classified as Firmicutes. The next two most prevalent phyla were Proteobacteria (12.5%) and Bacteroidetes (12.2%), however these phyla were overrepresented in two samples, biasing the total relative values. Deferribacteres were detected in only one sample (Koala 70, wet bottom present) and Acidobacteria were only detected in two (one clinically normal koala and one displaying wet bottom). Armatimonadetes was detected in three koalas without wet bottom, but in none of the five diseased koalas. These three phyla were detected at the lowest relative abundance across the ten samples. Data for relative read abundance for OTUs that could be taxonomically assigned to a genus level and occurred at a percentage of 0.01% or more in either group can be found in Table 2. This shows that the order *Lactobacillales*, and within that the genus *Aerococcus*, had the highest proportion of relative reads.

**Table 2.**
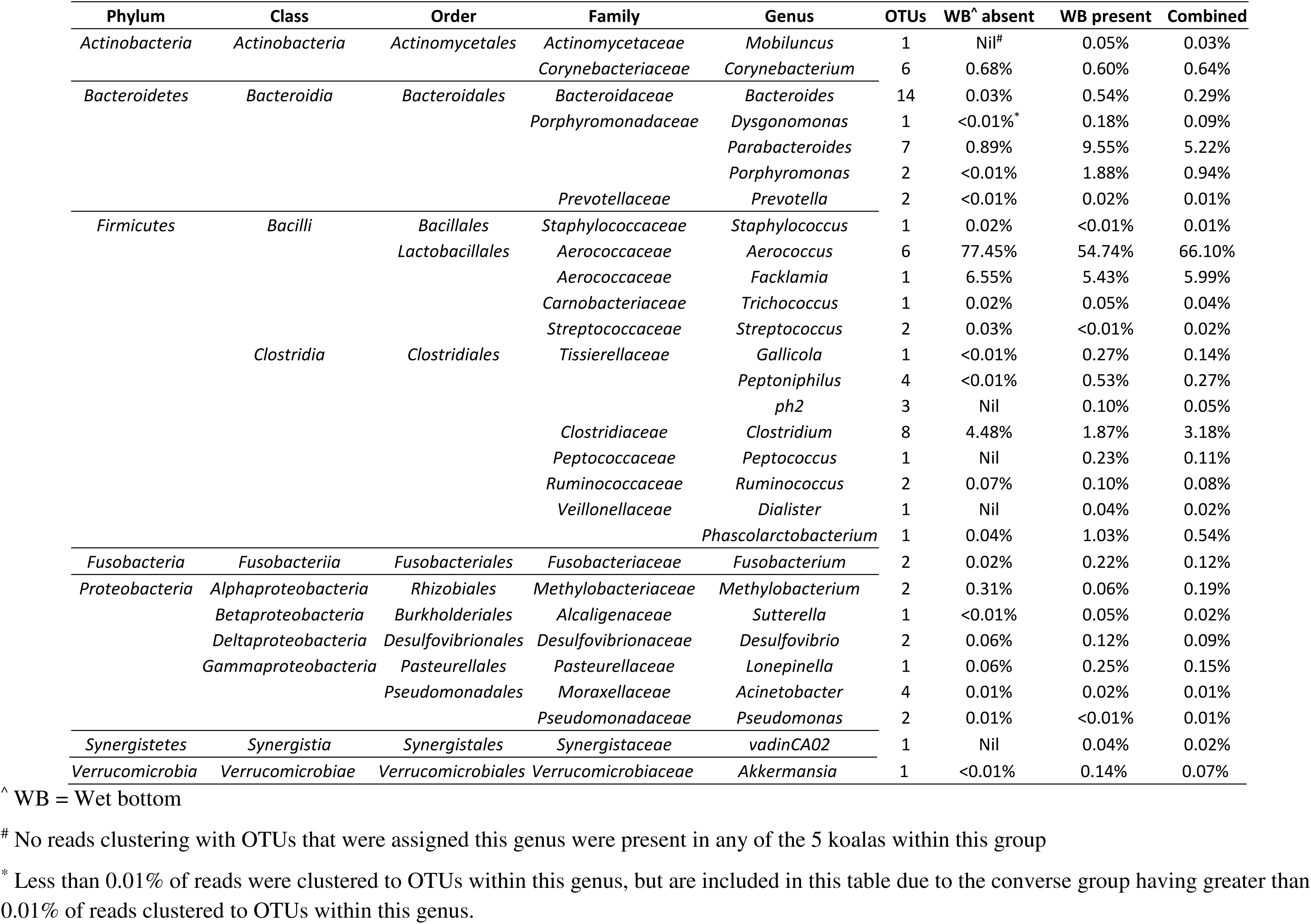
Relative abundance of OTUs with taxonomic classification to a genus level, in koalas with and without wet bottom. Only OTUs with relative abundance greater than 0.01% in at least one group are shown.

### Richness and diversity

Species richness within each sample is described in Table 1. The mean species richness and Chao1 from 100 iterations of subsampling every 5000 reads is shown in Figure 1. After 100 iterations of rarefaction to a depth of 160,000 reads per sample, the mean number of OTUs in the two groups was 80.0 (S.D. ± 9.62) and 75.93 (S.D. ± 24.61) for koalas with wet bottom and without wet bottom, respectively. All alpha diversity metrics compared between samples from koalas with or without wet bottom were not significantly different. This included observed OTUs (t = -0.31, *P* = 0.81), Chao1 (with wet bottom group (WB) mean = 90.7, without wet bottom group (NWB) mean = 88.4, t = -0.20, *P* = 0.83), phylogenetic diversity (WB mean = 7.8, NWB mean = 8.1, t = -0.39, *P* = 0.71) and Shannon’s diversity (WB mean = 2.4, NWB mean = 2.5, t = -0.15, *P* = 0.86) (see Table 3 for individual alpha diversity values). Results detailing abundance for all OTUs detected in koala urogenital samples are recorded in supplemental material Table S1.

**Figure 1.**
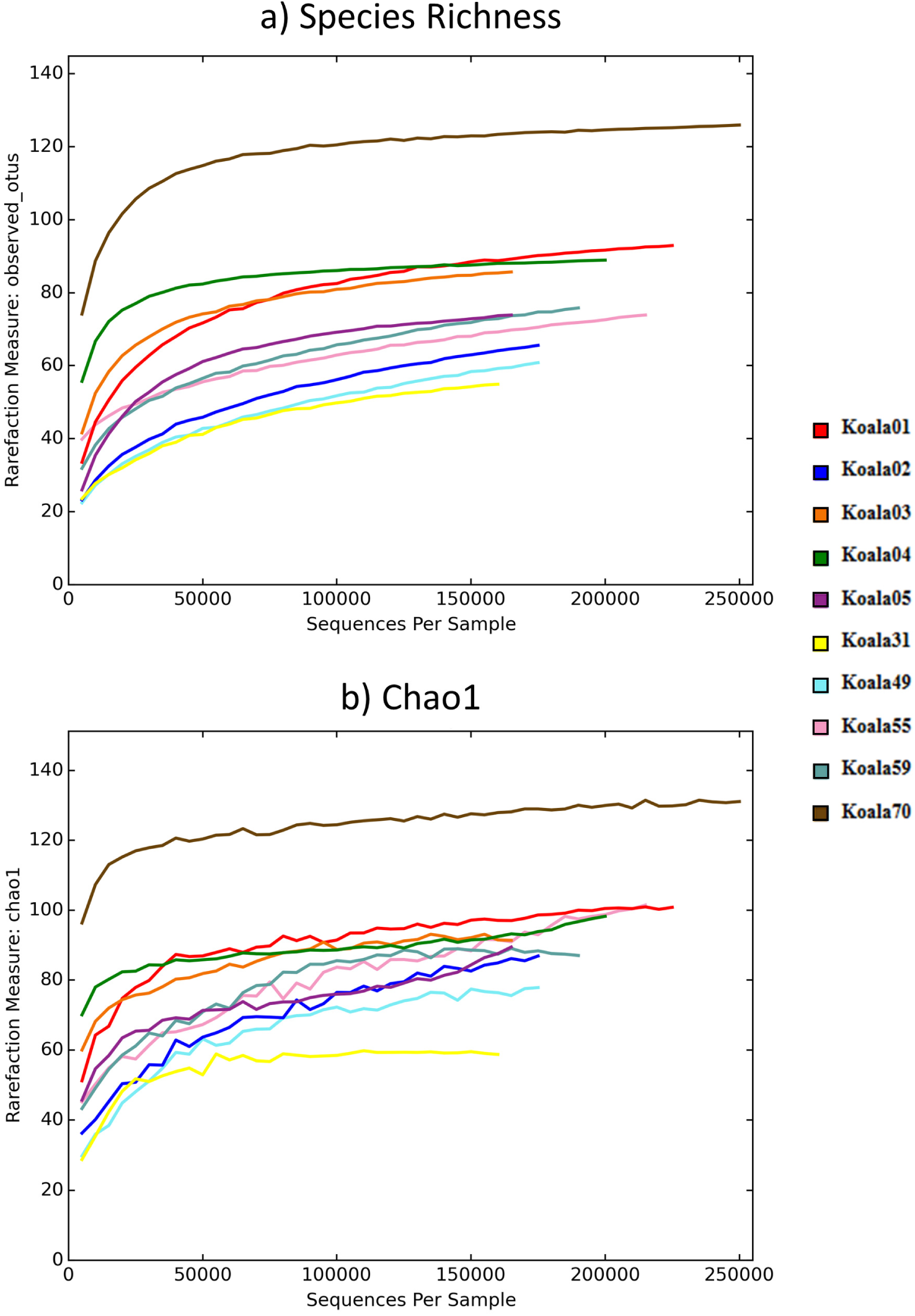
Rarefaction plots showing a) species richness (OTU abundance) and b) Chao1. OTUs were subsampled every 5000 reads, with 100 iterations, with the mean result of these iterations forming the plots. Koalas 1 – 5 were clinically normal (wet bottom absent), whilst koalas 31 – 70 had wet bottom.

**Table 3.** Alpha diversity metrics for microbial communities in the urogenital tract of koalas with and without wet bottom. All metrics assessed based on OTU values after subsampling to a depth of 160,000 reads, with 100 permutations. *P* values are non-parametric t-tests using 10,000 Monte Carlo permutations

|  | Richness (OTUs) | Shannon's diversity | Chao1 | Phylogenetic diversity |
| --- | --- | --- | --- | --- |
| Wet bottom absent |  |  |  |  |
| Koala 1 | 88.8 (± 1.7) * | 2.6 (± <0.01) | 97.1 (± 5.9) | 9.1 (± 0.2) |
| Koala 2 | 64.1 (± 1.2) | 2.7 (± <0.01) | 84.9 (± 7.4) | 7.0 (± 0.1) |
| Koala 3 | 85.4 (± 0.7) | 3.0 (± <0.01) | 91.5 (± 2.7) | 8.9 (± 0.1) |
| Koala 4 | 88 (± 0.9) | 3.1 (± <0.01) | 92.5 (± 3.7) | 7.7 (± 0.1) |
| Koala 5 | 73.7 (± 0.6) | 1.1 (± <0.01) | 87.6 (± 4.9) | 7.9 (± 0.1) |
| **Mean** | 80.0 (± 9.6) | 2.5 (± 0.7) | 90.7 (± 4.2) | 8.1 (± 0.8) |
| Wet bottom present |  |  |  |  |
| Koala 31 | 54.9 (± 0.3) | 2.4 (± <0.01) | 58.7 (± 0.8) | 6.5 (± 0.0) |
| Koala 49 | 59.2 (± 1.4) | 1.4 (± <0.01) | 76.4 (± 7.2) | 6.5 (± 0.2) |
| Koala 55 | 69.2 (± 1.9) | 2.3 (± <0.01) | 91.5 (± 13.5) | 7.8 (± 0.2) |
| Koala 59 | 72.9 (± 1.5) | 1.8 (± <0.01) | 87.4 (± 7.1) | 7.8 (± 0.1) |
| Koala 70 | 123.4 (± 1.3) | 4.1 (± <0.01) | 127.9 (± 5.9) | 10.4 (± 0.1) |
| **Mean** | 75.9 (± 24.6) | 2.4 (± 0.9) | 88.4 (± 22.8) | 7.8 (± 1.4) |
| *t* stat | -0.31 | -0.15 | -0.20 | -0.39 |
| *P* value | 0.81 | 0.86 | 0.83 | 0.71 |

\* All ± values are standard deviation from the mean

Fewer than half of the OTUs detected across the two sample groups were shared between them (112/254) (Figure 2). At a read depth of 160,000 there was a significant difference between the microbial communities in koalas with wet bottom compared to those without, based on the results of a 10,000 permutation PERMANOVA using Bray-Curtis dissimilarity (F = 4.92, *P* = 0.019) and unweighted (qualitative) UniFrac distances (F = 1.62, *P* = 0.031). There was no significant difference detected when using weighted (quantitative) UniFrac distances (F = 1.51, *P* = 0.061). 2D and 3D principal coordinate analysis (PCoA) graphs comparing koalas with and without wet bottom are shown in Figure 3. These identify two outliers in the wet bottom present group, koalas 49 and 70.

**Figure 2.**
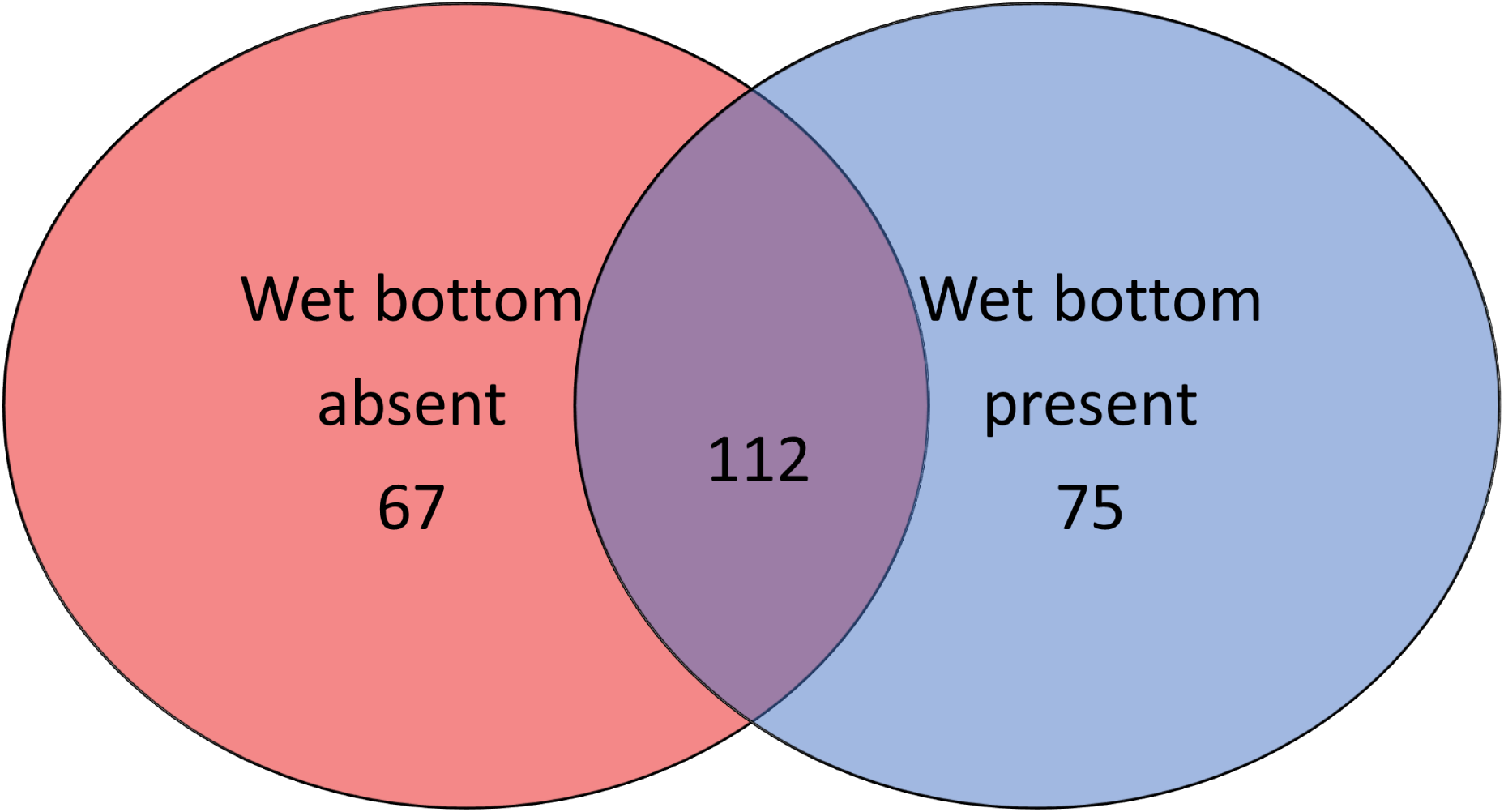
Venn diagram of the total operational taxonomic units (OTUs) detected in koalas with or without wet bottom. Overlap does not scale with OTU number.

**Figure 3.**
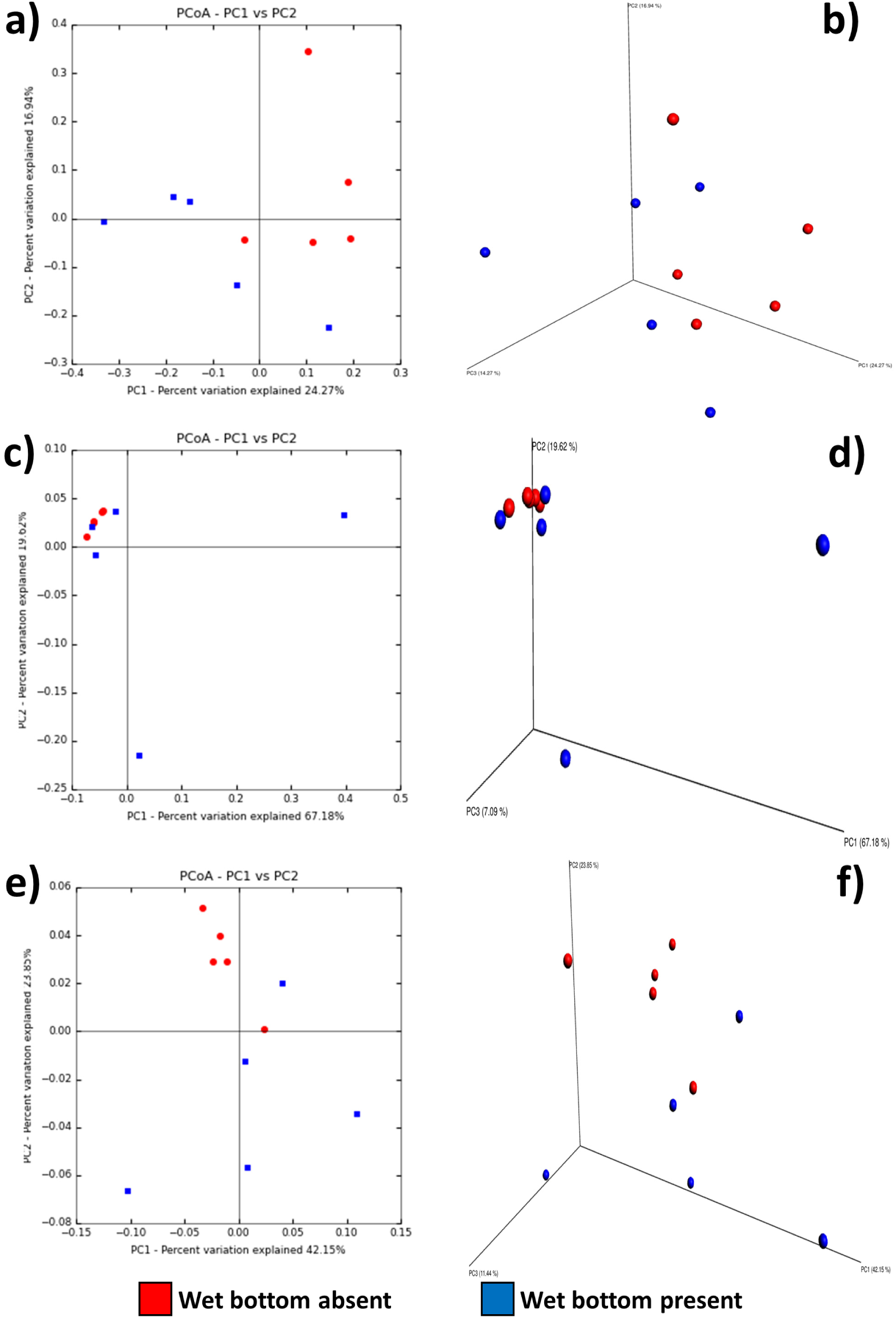
2D and 3D PCoA plots of koala samples, with and without wet bottom, using **a/b**) unweighted UniFrac distances of OTUs at a depth of 160,000 reads, **c/d**) weighted UniFrac distances of OTUs at a depth of 160,000, **e/f**) weighted UniFrac distances of normalised reads

### Comparisons between samples using DESeq2 normalised reads

Negative binomial normalisation of reads from each sample using DESeq2 still resulted in Firmicutes as the most dominant phylum across all samples. This was followed by Proteobacteria and Bacteroidetes (Figure 4). Overall there were 25 OTUs with significant (Benjamini and Hochberg (9) adjusted *P* < 0.05) over-representation or under-representation in wet bottom affected koalas, in comparison to clinically normal koalas, based on these normalised read counts (Table 4). Of those OTUs, when assessing absolute read count, six occurred only in koalas with wet bottom, whilst eight occurred only in koalas without wet bottom (Table 4). All normalised read values can be found in supplemental material Table S2, and all statistical comparisons in supplemental material Table S3.

**Figure 4.**
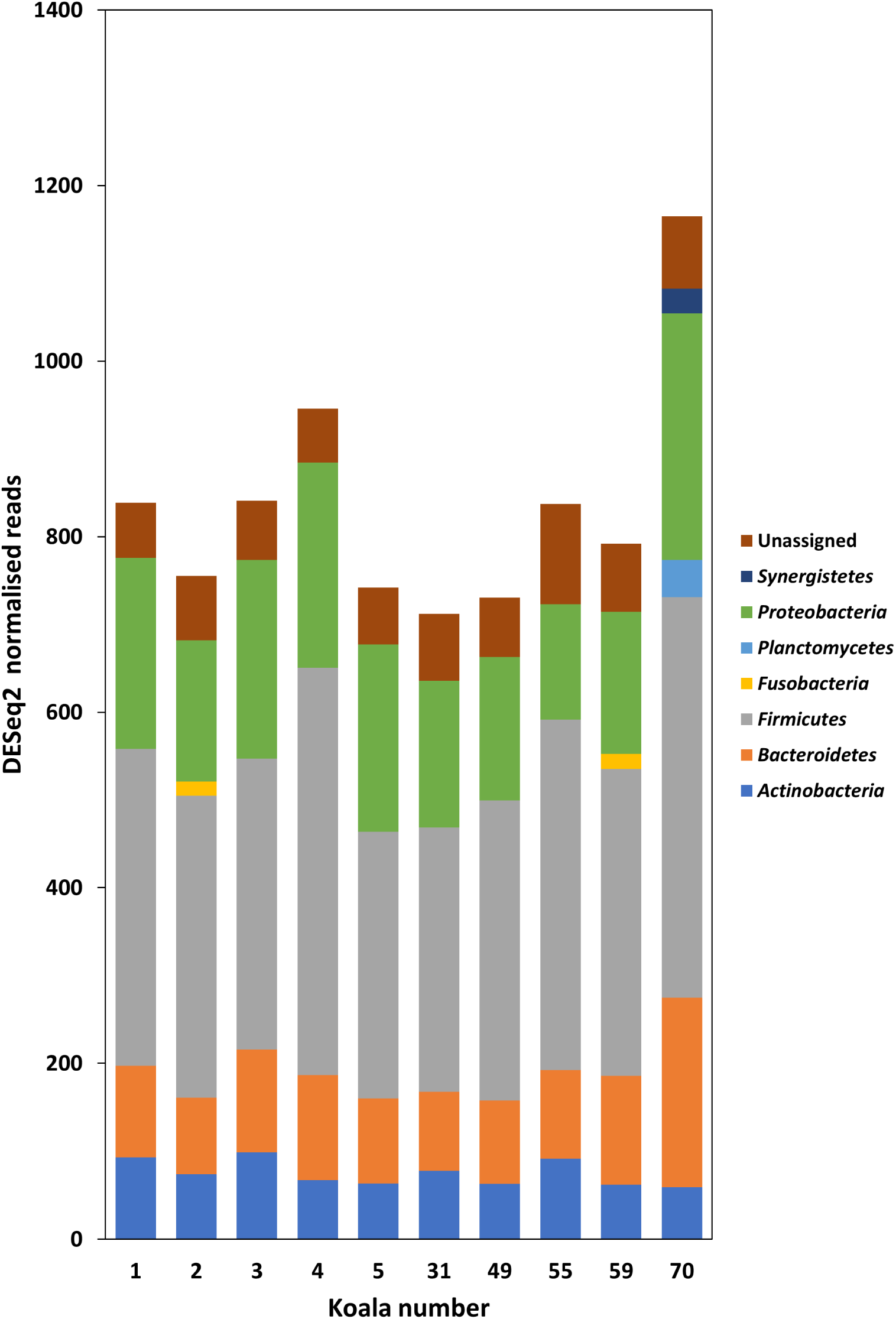
DESeq2 normalised read counts of phyla detected in koala urogenital swab samples. Phyla with fewer than 2% relative reads within each sample have been excluded for clarity. Reads were characterised into taxanomic groups using QIIME (40), utilising Greengenes (42) as a reference database. Koalas 1 – 5 were clinically normal (wet bottom absent), whilst koalas 31 – 70 had wet bottom

**Table 4.** Significant operational taxonomic units (OTU) assessed using DESeq2 (43), ordered from lowest to highest adjusted *P* value. Representative sequences were compared to NCBI nucleotide database using MegaBLAST (53), excluding ‘uncultured organisms’

| OTU ID | Adjusted *P* value<sup>#</sup> | Higher abundance group<sup>*</sup> | OTU present in samples/n |  | NCBI Mega BLAST <sup>^</sup> | Nucleotide Identity (%)<sup>^</sup> | Accession number<sup>^</sup> |
| --- | --- | --- | --- | --- | --- | --- | --- |
|  |  |  | WB absent | WB present |  |  |  |
| 38 | < 0.001 | WB present | 0/5 | 5/5 | *Peptoniphilus indolicus* | 96.8 | NR_117566 |
| 21 | < 0.001 | WB present | 1/5 | 5/5 | *Peptoniphilus asaccharolyticus* | 100 | KP944181 |
| 47 | < 0.001 | WB present | 0/5 | 3/5 | *Levyella massiliensis* | 100 | NR_133039 |
| 51 | < 0.001 | WB present | 0/5 | 3/5 | *Peptoniphilus lacrimalis* | 100 | KM624632 |
| 65 | 0.001 | WB present | 1/5 | 2/5 | *Sutterellaceae bacterium* | 99.5 | LK054638 |
| 86 | 0.003 | WB absent | 3/5 | 0/5 | *Bacteroides thetaiotaomicron* | 100 | KU234409 |
| 75 | 0.004 | WB absent | 2/5 | 0/5 | *Clostridium* sp. | 96.5 | AB622820 |
| 4 | 0.004 | WB absent | 5/5 | 5/5 | *Lactobacillales bacterium* | 92.8 | HQ115584 |
| 70 | 0.005 | WB absent | 2/5 | 0/5 | *Clostridium neopropionicum* | 94.6 | JQ897394 |
| 73 | 0.005 | WB present | 0/5 | 2/5 | *Alistipes onderdonkii* | 93.6 | NR_113151 |
| 69 | 0.005 | WB absent | 2/5 | 0/5 | *Lachnospiraceae bacterium* | 95.3 | EU728729 |
| 2 | 0.006 | WB absent | 5/5 | 5/5 | *Trichococcus* sp. | 94.2 | KU533824 |
| 94 | 0.007 | WB absent | 2/5 | 1/5 | *Rhizobiales* sp. | 100 | KJ016001 |
| 95 | 0.013 | WB absent | 2/5 | 0/5 | *Rhizobium leguminosarum* | 100 | KX346599 |
| 103 | 0.019 | WB absent | 2/5 | 0/5 | *Piscinibacter aquaticus* | 88.6 | NR_114061 |
| 106 | 0.019 | WB absent | 3/5 | 0/5 | *Burkholderia cenocepacia* | 100 | KU749979 |
| 109 | 0.019 | WB present | 0/5 | 2/5 | *Peptostreptococcus anaerobius* | 94.1 | NR_042847 |
| 148 | 0.019 | WB present | 0/5 | 2/5 | *Trichococcus* sp. | 87.5 | KU533824 |
| 159 | 0.019 | WB present | 2/5 | 4/5 | *Abiotrophia defectiva* | 87.9 | JF803600 |
| 114 | 0.019 | WB absent | 2/5 | 1/5 | *Massilia* sp. | 99.8 | JF279920 |
| 113 | 0.019 | WB absent | 3/5 | 0/5 | *Agrobacterium tumefaciens* | 100 | KU955329 |
| 1 | 0.030 | WB present | 5/5 | 5/5 | *Aerococcus viridans* | 95.1 | KC699123 |
| 105 | 0.035 | WB present | 4/5 | 5/5 | *Aerococcus sanguinicola* | 93.0 | LC145565 |
| 250 | 0.038 | WB present | 1/5 | 2/5 | *Hippea* sp. | 79.5 | FR754504 |
| 90 | 0.038 | WB present | 1/5 | 2/5 | *Olsenella scatoligenes* | 97.8 | NR_134781 |

## Discussion

Previous assessment of the koala microbiome has focused on the digestive system of koalas comparing either two free ranging animals from northern populations (10) or two captive koalas in Europe (11), from which the ocular microbiome was also assessed. This study is the first investigation of the microbiome of the urogenital tract of the female koala using modern high-throughput techniques, and only the second to assess the urogenital tract of a marsupial, with the tammar wallaby (*Macropus eugenii*) investigated previously using terminal restriction fragment length polymorphism analysis (12). The majority of reads in our sample set were classified in the order *Lactobacillales* (72.1%). This dominance of Firmicutes mirrors what has been seen in the human vaginal microbiome (13). In humans, the acidic pH of the genital tract is maintained by these lactic acid producing bacteria, which in turn is thought to play a role in preventing pathogenic infection (14). It appears from our sample set that koalas have a different family within the *Lactobacillales*, possibly performing a similar role. The most common family within our classified OTUs, in terms of either relative or normalised read abundance, was *Aerococcaceae*, whilst in humans the *Lactobacilli* dominate the reproductive tract. Within the *Aerococcaceae*, the genera *Aerococcus* and *Facklamia* were both represented in the top four most abundant OTUs. For all four significantly differentially abundant *Aerococcus* spp. OTUs, the same OTU could be detected in at least 4/5 (80%) of the converse sample group in absolute reads. For example, OTU 4, an *Aerococcus* sp. occurred in all ten koala samples, but was present in significantly higher quantities in clinically normal koalas after normalisation (*P* = 0.004). Whether specific *Aerococcus* spp. that are over or under-represented are an important factor in terms of disease presence requires further investigation. The production of hydrogen peroxide by commensal *Lactobacillus* species is thought to play a role in reducing the successful establishment of sexually transmitted diseases in humans (15, 16), and it has been shown that *Aerococcus* spp. are involved in hydrogen peroxide production (17, 18). In humans *Aerococcus* spp. have also been associated with disease, such as *Aerococcus urinae*, which can cause urinary tract infections (19) and septicaemia (20). Investigations into the urinary microbiome of women with and without ‘urgency urinary incontinence’ found that *Aerococcus* spp. were detected more frequently in cases where disease was present (21). In our study, the four *Aerococcus* spp. OTUs that had significantly different normalised abundance were evenly split, with two having higher abundance in koalas with wet bottom and two in koalas without wet bottom. The role of organisms within this family as opportunistic pathogens in koalas cannot be ruled out.

The *Aerococcus* were the most common genus amongst those OTUs with significant differential abundance after normalisation using DESeq2. The representative sequences of these four OTUs did not match known species within the *Aerococcus* genus, using the Greengenes database, with an identity greater than 90%, suggesting that these may represent novel species. This is not unexpected, as the culture of organisms from the koala urogenital tract has been limited to only a small number of studies, with the majority focused on diagnosing what was later deemed to be chlamydial infection (22-24). Efforts in culturing novel bacteria from koalas have focused primarily on its gut microbiome (25), of interest due to the koala’s unusual diet of *Eucalyptus* leaves, as well as the microbial flora in the pouch (26). Now that the we have identified (to the genus level) some organisms of interest in the female koala urogenital microbiome, it would be beneficial to use traditional microbiology techniques to further study these organisms. The other family of interest are the *Tissierellaceae*, within the order *Clostridiales*. The four *Tissierellaceae* OTUs with a significant differential abundance, all occurred in higher normalised quantities in koalas with wet bottom present. Three of these OTUs were in the genus *Peptoniphilus.* Interestingly, only one of these four OTUs was detected at all in the group of koalas without wet bottom, and only from the reads of one koala within this group. The *Peptoniphilus*, previously part of the genus *Peptostreptococcus* (27) within the family *Peptostreptococcaceae* (also in the order *Clostridiales*), have been associated with inflammatory diseases in other species. This includes mastitis in cattle (28) and pelvic inflammatory disease in humans (29). Organisms in this genus are fastidious anaerobes (27) and therefore potentially overlooked in culture based methods of investigating urogenital tract pathogens.

The average number of OTUs detected in our samples is difficult to compare to other publications investigating koala microbiomes. This is both due to the impact that sample site differences would have on OTUs present, as well as the method of OTU classification used. For instance, previous research on the koala intestinal microbiome used QIIME for analysis of 454 pyrosequencing reads (10) and detected 1855 OTUs, after removal of chimeras and singletons, from caecum, colon, and faecal samples. Similarly, an Illumina based study of microbiomes from ocular, oral, rectal and faecal samples from two captive koalas found OTU numbers ranging between 597 to 3,592, with a median of 1,456 (11). The average raw read numbers per sample assessed in these projects ranged from 12,831 (454 pyrosequencing) to 323,030 (Illumina). Our own average raw reads per sample were within that range (229,561), suggesting that the OTU differences between our studies are either associated with the sample site (urogenital versus digestive tract) or clustering methodology used. We employed UPARSE due to its demonstrated ability to correctly identify OTUs in a mock community and minimise spurious OTUs (30). Whilst there did not appear to be any strong clustering on our 2D or 3D PCoA plots, comparisons of the beta-diversity between groups highlighted that the makeup of the communities was significantly different when assessing both Bray-Curtis dissimilarity and unweighted UniFrac distances. These metrics assess presence/absence of OTUs between groups, with UniFrac also considering phylogenetic distance between OTUs present. Weighted UniFrac distances, which considers the abundance of individual OTUs, were not significantly different between groups. Therefore, koalas with and without wet bottom appear to have a significant difference in which OTUs are present in the samples, but not necessarily the abundance of OTUs between samples. Two samples had widely different OTU profiles (koala 49 and 70). This finding may support the hypothesis that wet bottom in female koalas without *C. pecorum* may be caused by more than one aetiological agent (5, 31). Further investigations to examine this hypothesis are indicated but require access to a large number of appropriately collected and stored samples. Such sample sets are currently not available for this species.

It could be argued that the skewed relative abundance of Proteobacteria and Bacteroidetes in the samples from koala 49 and 70, respectively, could be a result of swab contamination with faecal material, which would impact diversity inferences. The human microbiome project identified that reads from stool samples were predominately from the Bacteroidetes phylum (32), and the most recent assessment of the koala rectal microbiome found these two phyla to be the most abundant in samples taken from both koalas assessed (11). In koalas, the urogenital tract is accessed through the cloaca, which also contains the rectal opening. This makes faecal contamination difficult to avoid during sample collection. Future studies of the urogenital tract microbiome would benefit from either taking control samples from the rectum of the koala being sampled, or inverting the cloaca so that the urogenital opening is more easily accessible, as described previously for the tammar wallaby (12). In that study, approximately a quarter of phylotypes (26/96) were detected in both the urogenital and rectal samples, suggesting that bacteria being detected at multiple sites in marsupials is not unusual.

Our sample size is larger than previous studies of koala microbiomes, which have incorporated at most two individuals, yet it is substantially smaller than many studies in human medicine which include hundreds of samples (33). Our samples were opportunistically collected during population management exercises, and chosen from our sample archive due to the absence of *C. pecorum* from the French Island koala population at the time of testing (6). Whilst *C. pecorum* was subsequently determined to be present in this population (8), no koalas used in this project were positive via a *Chlamydiaceae* PCR. Importantly, no koalas used in this study were found to have reads classified within the *Chlamydiae* phylum after taxonomic assignment of OTUs.

Disturbance of the normal vaginal flora in humans, such as in cases of bacterial vaginosis, is a risk factor associated with infection by retroviruses (such as human immunodeficiency virus) and *Chlamydia trachomatis* (34). Our study provides useful data as to what bacteria could be expected in a clinically normal koala’s urogenital tract. This will allow for broader, more detailed studies on the impact that infection with *C. pecorum* has on the koala urogenital microbiome, and vice versa. Future studies would benefit from a greater sample size and a more diverse array of sampled regions both within a single state, and across the country. It would be interesting to follow the same individuals over time to determine if mating and breeding impact the microbiome of the urogenital tract, as occurs in humans (35). However, animal welfare issues regarding recapturing wild koalas multiple times may make this unfeasible. Additionally, as our study focused solely on female koalas, a follow up survey of the microbiome of the male urogenital tract would be enlightening. Finally, targeted studies assessing the prevalence of organisms associated with wet bottom would increase our understanding of organisms potentially impacting koala populations and could in turn assist with conservation of this iconic species.

## Methods

### Sample Collection and initial screening

Samples used in this study were urogenital swabs, from female koalas, stored in Buffer RLT (Qiagen) containing β-mercaptoethanol, taken from an archive of koala samples collected in 2011 from French Island, Victoria, Australia (38°21’0” S, 145°22’12” E). Koala samples were collected under general anaesthetic by veterinarians and trained field assistants during routine population management exercises and clinical health of koalas was recorded at the time. Sample collection was approved by the University of Melbourne Faculty of Veterinary Science Animal Ethics Committee, application ID: 1011687.1, and all sample collection was conducted following the Australian code for the care and use of animals for scientific purposes, 8th edition (36). Wet bottom score was assessed using a scoring system as previously described (37). These wet bottom scores grade the clinical findings relating to wet bottom from 0 (absent) to 10 (most severe). For the purpose of this study, koalas were grouped into wet bottom ‘present’ and wet bottom ‘absent’ categories. After screening all samples for *Chlamydiaceae* using a previously described qPCR (8, 38), we selected ten samples from female koalas where no *Chlamydiaceae* was detected. We used five samples collected from koalas showing no clinical signs of urogenital disease and five samples collected from koalas that showed clinical signs of wet bottom (Table 1).

### Amplification and sequencing

DNA extraction and amplification from the swab samples was performed commercially by The Australian Genome Research Facility (Australia). Variable regions three and four of bacterial 16S rRNA were amplified using primers 341F (5’ CCTAYGGGRBGCASCAG 3’) and 806R (5’ GGACTACNNGGGTATCTAAT 3’). Sequencing was performed on the Illumina MiSeq platform to produce paired end reads of 300 bp (2 × 300 bp).

### Quality filtering and OTU assignment

Quality filtering and operational taxonomic unit (OTU) assignment was undertaken using a mixture of scripts and algorithms available in the programs USEARCH 8.1 (39) and QIIME 1.9.1 (Quantitative Insights Into Microbial Ecology) (40). Unless otherwise stated, default settings were used for all scripts. Read processing to reduce errors was undertaken as described by Edgar and Flyvbjerg (41). The forward and reverse 300 bp paired-end reads for each swab sample were merged using the USEARCH script **fastq_mergepairs**. In this process, the Phred score of overlapping bases is recalculated to improve error calling. Bases with the same nucleotide called in both the forward and reverse reads have an increased recalculated score, and those with disagreements are reduced. This increases confidence in the calculated error probability of the merged reads. Primers were then trimmed from the 5’ and 3’ ends of the merged reads using seqtk (https://github.com/lh3/seqtk). Trimmed reads were filtered for quality using the USEARCH script **fastq_filter**. This script filters reads using the maximum expected errors per merged read. The number of expected errors is obtained by the sum of the Phred derived error probability. If the expected number of errors is less than one, then the most probable number of errors is zero (41). We utilised a maximum expected error threshold of 1, resulting in reads with an error probability of 1 or greater being removed. In addition to using the number of expected errors for filtering, trimmed reads shorter than 400 bp were discarded. Unique reads within the entire sample set were assigned OTUs using the USEARCH algorithms **derep_fulllength** and **cluster_otus** (30), with a minimum identity of 97% for clustering, or a cluster radius of 3.0. Chimeras are filtered from the sample set within the **cluster_otus** command using the UPARSE-REF maximum parsimony algorithm (30). Singletons were excluded from OTU clustering due to the high likelihood that they contain errors (30, 41). The merged/trimmed reads from each swab sample, including the previously excluded singletons were matched with the clustered OTUs using USEARCH script **usearch_global**, with a threshold of 97% identity to group a read into a specific OTU. The taxonomy of each OTU was determined by using the QIIME script **assign_taxonomy**.py in conjunction with the Greengenes taxonomy database (version 13_5, 97% clustered OTUs) (42). This script utilises the UCLUST algorithm (39) to identify a consensus taxonomy of the reads within an OTU against the curated database, based on a similarity of 90% and a minimum consensus fraction of 0.51. Chloroplast and mitochondrial OTUs were removed from the dataset using the QIIME script **filter_taxa_from_otu_table.py**.

### Read normalisation and analysis

Read data was assessed using three different methods. Relative abundance was utilised to compare basic phylum presence in each sample. Rarefaction of reads was undertaken, using **multiple_rarefactions.py** QIIME script, to assess alpha and beta diversity at a set read level. Negative-binomial normalisation of reads, using DESeq2 (43) as recommended by McMurdie and Holmes (44), was performed using the QIIME script **normalize_table.py.** For rarefactions, reads within each sample are subsampled (without replacement) every 5000 reads, from 5000 to 250,000 reads. This represented the maximum number of reads present in the sample with the most reads (rounded down to the nearest value divisible by 5,000). At each step, 100 permutations were undertaken. Alpha-diversity metrics (including species richness, Chao1 (45), phylogenetic diversity and Shannon’s diversity (46)) were generated for each step. Comparisons of these values were undertaken using values obtained after subsampling to a depth of 160,000. This equalled the sample with the fewest reads (rounded down to the nearest value divisible by 5,000). Non-parametric comparisons of mean alpha diversity metrics between the two sample groups (wet bottom present or absent) were undertaken with the **compare_alpha_diversity.py** QIIME script. This script utilised a non-parametric two sample t-test with 10,000 Monte Carlo permutations to determine whether the alpha diversity was significantly different between the two groups (wet bottom present/absent) at a depth of 160,000 reads. Beta-diversity was assessed at the same depth as above (160,000 reads) using the **beta_diversity_through_plots.py** QIIME script, in which both unweighted and weighted UniFrac distances (47) were assessed. Bray-Curtis dissimilarity (48) between samples was also assessed. The analysis of beta-diversity required a phylogenetic tree. For this, an alignment of representative sequences of each OTU was created with PyNAST (49) and UCLUST using the **align_seqs.py** QIIME script. A tree was produced from this alignment using FastTree (50), and used as input for beta-diversity analysis. **beta_diversity_through_plots.py** produced distance matrices for each of the tests (UniFrac and Bray-Curtis), from which principal coordinates and eigen values could be calculated. PCoA plots using the 2 or 3 most influential principal coordinates were drawn from the resulting distance matrices either using either the **make_2d_plots.py** QIIME script, or within the **beta_diversity_through_plots.py** script using EMPeror 9.51 software (51), respectively. Distance and dissimilarity metrics were used to compare the microbial communities between the two groups by utilising the permutational ANOVA (PERMANOVA) method within the **compare_categories.py** QIIME script, with 10,000 permutations. Statistical comparisons of the differential abundance of OTUs between koalas with and without wet bottom utilised DESeq2 within the QIIME script **differential_abundance.py.** These comparisons aimed to determine OTUs which were overrepresented in either group. Statistically significant results, using DESeq2s negative binomial Wald test, were based on *P*-values < 0.05, and were adjusted for false discovery within the script, using the method described by Benjamini and Hochberg (9).

The NCBI nucleotide database (52) was utilised to search for species level matches of significantly differentially abundant OTUs. This was conducted using the representative sequence of the significant OTU and the MegaBLAST algorithm (53), excluding uncultured sample sequences.

All reads used in the project are available through the NCBI BioProject ID: PRJNA359726. Accession numbers (SRX2464137 – SRX246146) for short reads are available in the short-read archive.

## Acknowledgements

Alistair Legione is supported by an Australian Postgraduate Award. Funding for the research described was provided by the Holsworth Wildlife Research Endowment – Equity Trustees. The authors declare that there are no competing financial interests in relation to this research.

The authors would like to acknowledge the guidance and advice of Mr. Brendan Ansell, as well as those who assisted with the collection of samples from koalas in the field.

